# Sexually antagonistic co-evolution can explain female display signals and male sensory adaptations

**DOI:** 10.1101/2022.03.14.484300

**Authors:** R. Axel W. Wiberg, Rosalind L. Murray, Elizabeth Herridge, Varpu Pärssinen, Darryl T. Gwynne, Luc F. Bussière

**Affiliations:** Biological and Environmental Sciences, University of Stirling, Stirling, FK9 4LA, United Kingdom; Department of Biology, University of Toronto at Mississauga, Ontario, L5L 1C6, Canada; Department of Zoology, Stockholm University, Svante Arrhenius väg 18B, Stockholm, Sweden, 106 91; Health and Social Care Analytical Services, St Andrews House, Regent Road, Edinburgh, EH1 3DG, United Kingdom; Institut för Biologi och Miljövetenskap, University of Gothenburg, Box 451, Göteborg, Sweden, 405 30; Gothenburg Global Biodiversity Centre, University of Gothenburg, Box 451, Göteborg, Sweden, 405 30

## Abstract

The prevalence and diversity of female ornaments poses a challenge to evolutionary theory because males should prefer mates that spend resources on offspring rather than ornaments. Among dance flies, there is extraordinary variation in sexual dimorphism. Females of many species have conspicuous ornaments (leg scales and inflatable abdominal sacs). Meanwhile males of some species have exaggerated regions of their eyes with larger ommatidial facets that allow for regionally elevated photosensitivity and/or acuity. Here, we conduct a comparative study of these traits using both species descriptions available from the literature, as well as quantitative measures of eyes and ornaments from wild-caught flies. We show a conspicuous covariance across species between exaggerated male dorsal eye regions and the extent of female ornaments: species with highly ornamented females have males with more exaggerated eyes. We discuss this pattern in the context of competing hypotheses for the evolution of these traits and propose a plausible role for sexually antagonistic coevolution.

## Introduction

Female secondary sexual traits, such as ornaments, remain poorly understood in evolutionary biology. This is, in part, because female fecundity is often resource-limited, and males should avoid females that divert resources away from offspring towards extravagant morphologies (Fitzpatrick et al. 1995; Clutton-Brock 2009). Although some examples of such traits exist (Tobias et al. 2012; Dougherty 2021), and to greater extents in some clades (e.g. Kraaijeveld et al. 2007), female ornamentation remains generally rare in nature. Efforts to explain the evolution of female ornamentation have focused largely on their evolution as a result of male preferences for more ornamented females, whereby the ornaments are inter- or intra-sexual indicators of fecundity, fertility, or quality (Schlupp 2018; Fitzpatrick and Servedio 2018). This theoretical framing also informs much of the empirical work in this field (e.g. Weiss and Dubin 2018; Yong et al. 2018; Kopena et al. 2018; Higham et al. 2021; Nolazco et al. 2022). By contrast, theory developed for male-biased display traits allows for more varied processes including direct or indirect benefit signalling, but also exploitation of sensory biases, and sexual conflict (e.g. Sakaluk 2000; Macías Garcia and Ramirez 2005; Nakano et al. 2010; Kolm et al. 2012; Macías Garcia et al. 2012; Amcoff and Kolm 2015; Nakano et al. 2013).

The dance flies of the subfamily Empidinae (Diptera: Empididae) are a speciose group with a global distribution, and exhibit remarkable diversity in mating systems and sexual dimorphisms among both sexes (Collin 1961; Cumming 1994; Chvala 2005; Murray et al. 2022). The ancestral mating behaviour appears to have involved the aerial transfer of a nuptial gift (typically, an exogenous oral nuptial gift, *sensu* Lewis *et al.,* 2014) from the male upon which the female feeds during copulation (Downes 1970; Cumming 1994). Females generally do not hunt as adults (Newkirk 1970), and these protein-rich gifts are apparently necessary for egg development (Hunter & Bussière 2019). In many species, contests among females for access to nuptial gifts are so intense that the operational sex ratio (OSR; Emlen and Oring 1977; Ahnesjö *et al.,* 2002) in aerial mating swarms is heavily female-biased. Intense contests among females for nuptial gifts seem to have promoted the evolution of several secondary sexual characters in females (Cumming 1994), including enlarged or coloured wings, inflatable abdominal sacs and, most frequently, pinnate scales on one or more pairs of legs (see figure 1) which females often wrap around their abdomens in flight and serve to attract and secure a male with a nuptial gift (Murray *et al.,* 2018). There remains some debate around the extent to which these female characters are honest signals of condition or fecundity or rather represent attempts at deception and sexual conflict (Funk and Tallamy 2000; Browne and Gwynne 2022). If female ornaments are in fact sexually antagonistic deceptive signals, male resistance traits are expected to evolve in some well-adorned species (Arnqvist and Rowe 2005).

**Figure 1.**
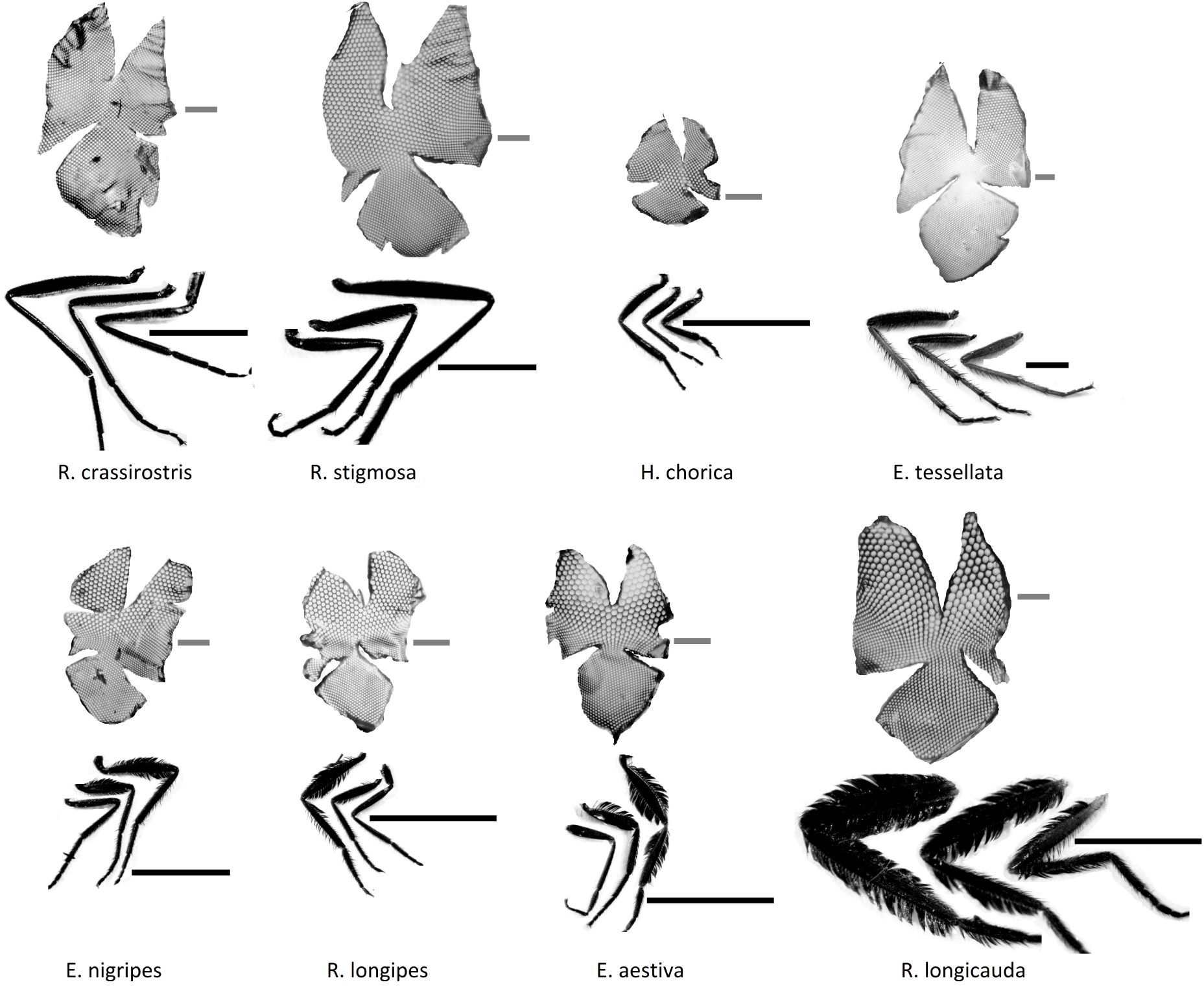
Micrographs of nail varnish eye casts and female front, middle and hind legs. Each of the eight species included in the comparative analysis is represented. The placement of these species in the new phylogenetic tree is highlighted in figure S1. Images are taken at different magnifications; black scale bars to the right of the respective female legs represent 1 mm, while grey scale bars to the right of the respective eye casts represent 0.1 mm.

Meanwhile, males of some dance fly species have dichoptic compound eyes, with dorsal eye regions showing enlarged ommatidia in comparison to females (Collin 1961; see figure 1). These areas can enhance certain visual capacities including photosensitivity (referred to as “bright zones”), or, when paired with a less curved eye surface, improve visual acuity (“acute zones”; Land 1989; 1997). These adaptations can locally fine-tune optimal visual capabilities, but come at the cost of visual ability in other regions of the eye. Such trade-offs are due to constraints imposed on the resolving power of the compound eye by an upper limit on eye size (Land 1989; 1997). Previous hypotheses to explain variation in male eye morphology have explicitly or implicitly invoked selection among males for faster mate acquisition or choice of more ornamented females (Downes 1970; Thornhill and Alcock 1983). For example, selection might favour specialised eyes as adaptations to “scramble-like” male competition as in other systems (e.g. Eichorn *et al.,* 2017). Alternatively, they have been proposed as adaptations to assess honest female ornaments (LeBas et al., 2003). An unexplored hypothesis is that these adaptations, that presumably augment visual perception and acuity, are exactly the sorts of resistance traits that are expected if female ornaments are generally deceptive and sexually antagonistic (Arnqvist and Rowe 2005).

Although a link between male eye morphology and female ornamentation in dance flies has been proposed (Downes 1970; Thornhill and Alcock 1983; LeBas et al., 2003), the degree of variation in eye dimorphism and whether these traits covary across species has not been investigated. Here we conduct a comparative study investigating the co-evolution of female ornaments and male eye morphology in dance flies. We first make use of a survey of species descriptions on morphology to assess broad patterns of association between eye dimorphism and ornament dimorphism among many dance flies. We then quantify associations between continuous measurements of male and female eye morphology and the degree of female ornamentation for a subset of species. If female ornaments and male eye exaggerations represent a sexually antagonistic arms race, we expect to observe a positive association between the two traits. We discuss our findings in the context of competing hypotheses that link the evolution of female ornamentation and male eye adaptations in these species.

## Methods

### Data collection and field sampling

We examined species descriptions in Collin (1961) to document possible associations between female ornaments and eye sexual dimorphism among the British dance fly species of the subfamily Empidinae. Our use of descriptions in this key reflects both our focus on local fauna but also a bias resulting from the fact that comprehensive descriptions are largely only available for species from Northern Europe and the British Isles. However, we note that there is extraordinary diversity from other geographic regions (e.g. Daugeron et al. 2011), and our results must be interpreted with this in mind. We noted the presence or absence of four main categories of sexually dimorphic traits in females that are likely to be sexually selected ornaments: enlarged wing size, darker wing colouration, pinnate scales on legs, and inflatable abdominal sacs. We also noted any descriptions of sexual dimorphism in eye facets. We collected information for all 95 species in the genera *Empis* and *Rhamphomyia*, and filtered the data to 93 species for which we could obtain information of female leg pinnation and/or other ornamentation, and descriptions of male eye morphology. We included information from one outgroup in the genus *Hilara* for phylogenetically corrected analyses (see below).

In order to assess the covariance between continuous measures of ornaments and eye morphology, we collected dance fly samples during the summers 2009, 2010 and 2011 on the eastern side of Loch Lomond between the Scottish Centre for Ecology and the Natural Environment (SCENE) and Rowardennan, Scotland, and during the summer of 2012 near Glen Williams in Ontario, Canada. We caught flies from mating swarms or when resting on vegetation using sweep nets (Murray *et al.,* 2017). Of the sampled taxa, we selected eight species, *R. longicauda*, *R. longipes*, *R. stigmosa*, *R. crassirostris*, *E. aestiva*, *E. tessellata*, *E. nigripes*, and *H. chorica*, that represent a range of variation in female ornaments, male eye morphologies, and mating systems (e.g. OSR, type of nuptial gift; see also figure 1 and table S2 for final sample sizes).

### Morphological Measures

Differences in eye morphology were quantified from nail varnish casts (see figure 1; Ribi *et al.,* 1986; Narendra *et al.,* 2010). Varnish was painted onto the eyes of specimens, allowed to dry, and then carefully removed. Three small cuts (figure 1) were then made to flatten the varnish casts and allow micrography for digital measurements. The diameters of seven neighbouring ommatidia on the dorsal and ventral portions of the eyes were measured from micrographs using tpsUtil (v 1.58; Rholf 2017) and tpsDig2 (v 2.17; Rholf 2017) in a manner similar to Döring and Spaethe (2009). We selected ommatidia located as dorsally or as ventrally as possible while accounting for cutting locations, obvious distortions in the nail varnish casts, and ensuring entire units of one central ommatidia and six surrounding ommatidia were measured. The difference between mean dorsal and ventral diameters was standardised by thorax length and taken as a measure of exaggeration. Thus, a positive value of exaggeration indicates larger dorsal ommatidia and a negative value indicates larger ventral ommatidia.

Female ornamentation was quantified by first computing the total leg area of the four posterior legs of females for each species using the software ImageJ (v 1.48; Rasband 2016). While these ornament indices underestimate ornament expression for some species (*e.g.,* for species where females have multiple ornaments, such as inflated abdominal sacs and/or darker wings in addition to pinnate scales), they nevertheless allowed us to measure trait expression in an analogous way across multiple taxa. Because the most heavily adorned species also had elaborate pinnate scales, ignoring the other ornament types makes our assessment of covariance between eyes and ornament expression conservative. Moreover, our results from the literature search of Collin’s (1961) key indicate that pinnation itself may be an especially strong predictor (see Results below). These measures were scaled by the total length of the femora and tibia of all four legs. Finally, the mean scaled male leg area was subtracted from each individual female measure for each species to give individual measures of ornamentation for each female.

### Sequence data collection and phylogenetics

We obtained sequence data for the carbamoylphosphate synthase domain of the rudimentary locus (CAD) and the mitochondrial locus *Cytochrome oxidase I* (COI) from the literature (Watts et al. 2016; Wahlberg & Johanson 2018; Murray et al. 2020), and from the BOLD database (Ratnasingham et al. 2007; https://v3.boldsystems.org). After curating the data to a single sequence per species, favouring longer sequences with fewer gaps for each species, the final dataset comprised 62 species from the genera *Empis*, *Rhamphomyia*, and *Hilara* as well as a further outgroup *Heterophlebus versabilis*, used also to root the tree in Murray et al. 2020 (see table S3 for sequences and sources).

We then aligned the sequences for each locus separately with MAFFT (v. 7.505; Katoh et al. 2013) and concatenated the alignments. Note that not all species had a representative sequence for both loci (table S3). Where this was the case, the species received a sequence of gap characters in the concatenated alignment. We then performed maximum likelihood tree construction with IQ-Tree (v. 2.0.7; Nguyen et al. 2015) setting different gene partitions for the two loci. We allowed IQ-Tree to test and pick the best substitution model (“-m TEST”). Comparing the subset of this new phylogeny to that of Murray et al. (2020) finds a Robinson-Fould’s distance of 0.29, indicating that the majority of the splits are in agreement. The structure of the new phylogenetic tree is shown in figure S1.

### Statistical analyses

We tested for an association between the categorical information on eye dimorphism and female ornamentation collected for a total of 93 species from Collin (1961) using several Chi-squared tests to evaluate whether ornamental traits were associated with eye dimorphism (figure S2-S4). The first test considered the presence of ornament, regardless of type (i.e. enlarged or darkened wings, inflatable abdominal sacs, pinnate leg scales). We also tested for association between eye exaggeration and either a) pinnate leg scales or b) all other ornaments excluding pinnate leg scales. This final contingency table has expected frequencies < 5 in some cells (figure S4), necessitating the use of a Fisher’s Exact Test.

For a subset of the data from the taxonomic descriptions, we could perform more robust testing using phylogenetically corrected mixed effects models using our new phylogeny (30 species, or 32% of the full data). We therefore pruned the tree to contain only these species, and fit a model with the presence/absence of male dorsal facet exaggerations as the response variable, and female pinnate scales as a predictor. Since male eye morphology was coded as 0 (no enlarged dorsal facets) or 1 (enlarged dorsal facets), we applied a Bernoulli error model (family=bernoulli(), using a logit link function).

For the eight species (*R. longicauda*, *R. longipes*, *R. stigmosa*, *R. crassirostris*, *E. aestiva*, *E. tessellata*, *E. nigripes*, and *H. chorica*) from which we obtained continuous morphological measures, the relationship between ornamentation, sex and eye exaggeration was also tested in a mixed model framework, fitting ornamentation, sex, and their interaction to predict eye exaggeration. The species studied here are from two genera, but recent molecular phylogenetic work shows that these genera are not monophyletic, and that female ornamental traits have evolved independently several times (Watts et al., 2016; Murray et al., 2020). To include phylogenetic information, we used the consensus tree from Murray et al., (2020), pruned to contain the eight species of interest. The results are the same whether we use the phylogeny of Murray et al. (2020) or the new one from this study. We report results using the phylogenetic tree from Murray et al. (2020) in this analysis because there is an associated posterior distribution of trees. Thus, in addition to the main analysis on the consensus tree, we also marginalised our analyses over a random sample of trees from the. Previous studies have suggested similar approaches to account for uncertainties in the topology of the tree (Hadfield and Nakagawa 2010; Hadfield 2010; Ross et al., 2013; Murray et al., 2020). Although simulations have suggested that as few as 50 trees from the posterior distribution may be required to adequately account for phylogenetic uncertainty (Nakagawa & de Villemereuil 2019), we sampled 200. We did this by fitting several models, each with a different input tree (with the function brm_multiple()), and merging posterior samples. In all models we also included a second random effect to account for multiple measures from each species. Continuous predictors were scaled (the mean subtracted from each value) prior to analysis (Schielzeth et al., 2010).

All analyses were performed in R (v.4.3.3; R Core Team 2024) using the packages “ape” (v. 5.4; Paradis et al., 2004), “brms” (v. 2.20.4; Bürkner 2017), “loo” (v. 2.7.0; Vehtari et al. 2024) and “coda” (v. 0.19.4; Plummer et al. 2006). Figures were plotted with “ggplot2” (v. 3.3.1; Wickham 2016). We follow Muff et al. (2022) and report results using the “language of evidence” for results of frequentist statistics and evaluate Bayesian results in terms of the entire posterior distributions and references to the 89% highest posterior density intervals (HPDIs; McElreath 2018).

## Results

We found strong evidence for positive associations between ornament expression and eye exaggeration from the 93 taxonomic descriptions of female ornaments and male eyes (figures S2-S4). Dimorphism in eye morphology was more strongly associated with pinnate leg scales than other ornament characters (figure S2, all ornaments together: Chi-squared = 11.63, d.f. = 1, p < 0.001; figure S3, pinnate scales only: Chi-squared = 17.02, d.f. = 1, p < 0.001; figure S4). In contrast, there was no evidence for an association with other ornaments excluding pinnate scales (Fisher’s Exact Test: Odds Ratio (95% CI) = 0.76 (0.18, 2.93), p = 0.77; figure S3).

These same patterns are also reflected in the phylogenetically corrected analysis of the subset of 30 species. Although the 89% highest posterior density interval (89% HPDI) for the effect of pinnation includes 0 (figure 2), 93% of the full posterior distribution lies above 0. The weight of the evidence thus indicates a decidedly positive association between female leg pinnation and male eye exaggerations. Importantly, all effects are in the same direction; species with female ornamentation or leg pinnation are more likely to also have exaggerated eye morphology (figures S2-S4 and figure 2). The intercept term estimated at below zero 0 suggests that species without descriptions of female pinnation have under a 50% chance of exhibiting exaggerated eye morphologies (figure 2). The posterior mean of phylogenetic heritability was 0.24. Although most of the posterior suggested relatively modest phylogenetic inertia, the 89% HPDI was extremely wide, with 5.4% of the distribution > 0.5 (figure S5). Moreover, Leave-One-Out (LOO) model comparisons indicate that a model without phylogenetic effects is a better fit to the data (ELPD_phylogeny_ = -21.4, ELPD_no-hylogeny_ = -20.2; SE_diff_ = 0.6).

**Figure 2.**
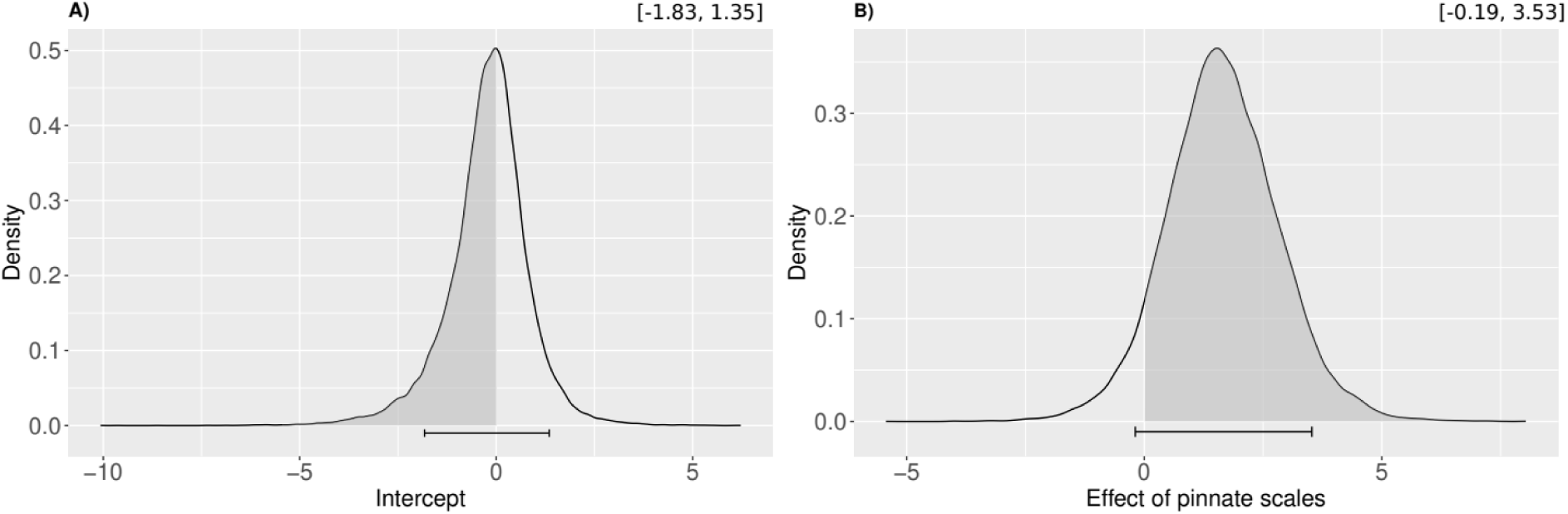
Posterior distributions of fixed effects from a phylogenetically controlled model of exaggerations of male dorsal acute zones as a function of presence or absence of female pinnate scales in Empidine dance flies. **A)** posterior for the intercept term and **B)** posterior for the effect of the species having pinnate scales. Shaded areas under the curve depict the proportions of the distribution < 0 (in panel A) or > 0 (in panel B). Horizontal error bars show the 89% HPD intervals. Inset text above each panel give the interval ranges to three significant figures. Trace plots of MCMC chains are shown in figure S6 along with traces of phylogenetic effects.

Within the subset of eight species for which we collected wild specimens, we also found strong evidence for an association between continuous measures of female ornamentation and eye exaggeration, with more exaggerated eyes occurring in taxa with more ornate females (figures 3 and 4). Posterior distributions for the sex by female ornamentation interaction term never overlapped 0, and the 89% HDPI ranged from 0.0921 to 0.108 (figure 3D). Intriguingly, female eyes in ornamented species tended to show the opposite pattern to males (89% HDPI = [-0.055, -0.0128]; figure 3B); females had larger ventral facets than dorsal ones, and more ornamented species tended to have more exaggerated ventral facets (partial slopes for females were negative; figures 3 and 4). Finally, the intercepts near 0, with a slight positive deviation for males, suggest that eyes of unornamented species have fairly uniform ommatidia sizes and are much less dimorphic, perhaps reflecting an ancestral condition (figures 3 and 4). The evidence for these patterns remained strong in models that controlled for phylogeny while accounting for phylogenetic uncertainty by marginalising over a sample of the posterior distribution of trees (figures S8 and S9). Posterior means of phylogenetic heritability were low to moderate (0.41-0.45), but again there was considerable uncertainty in these estimates for both models (figure S10). Indeed, LOO model comparison of the model including the consensus phylogenetic tree to a model without the tree only marginally preferred including the phylogeny (ELPD_phylogeny_ = 663.9, ELPD_no-hylogeny_ = 663.8; SE_diff_ = 0.2).

**Figure 3.**
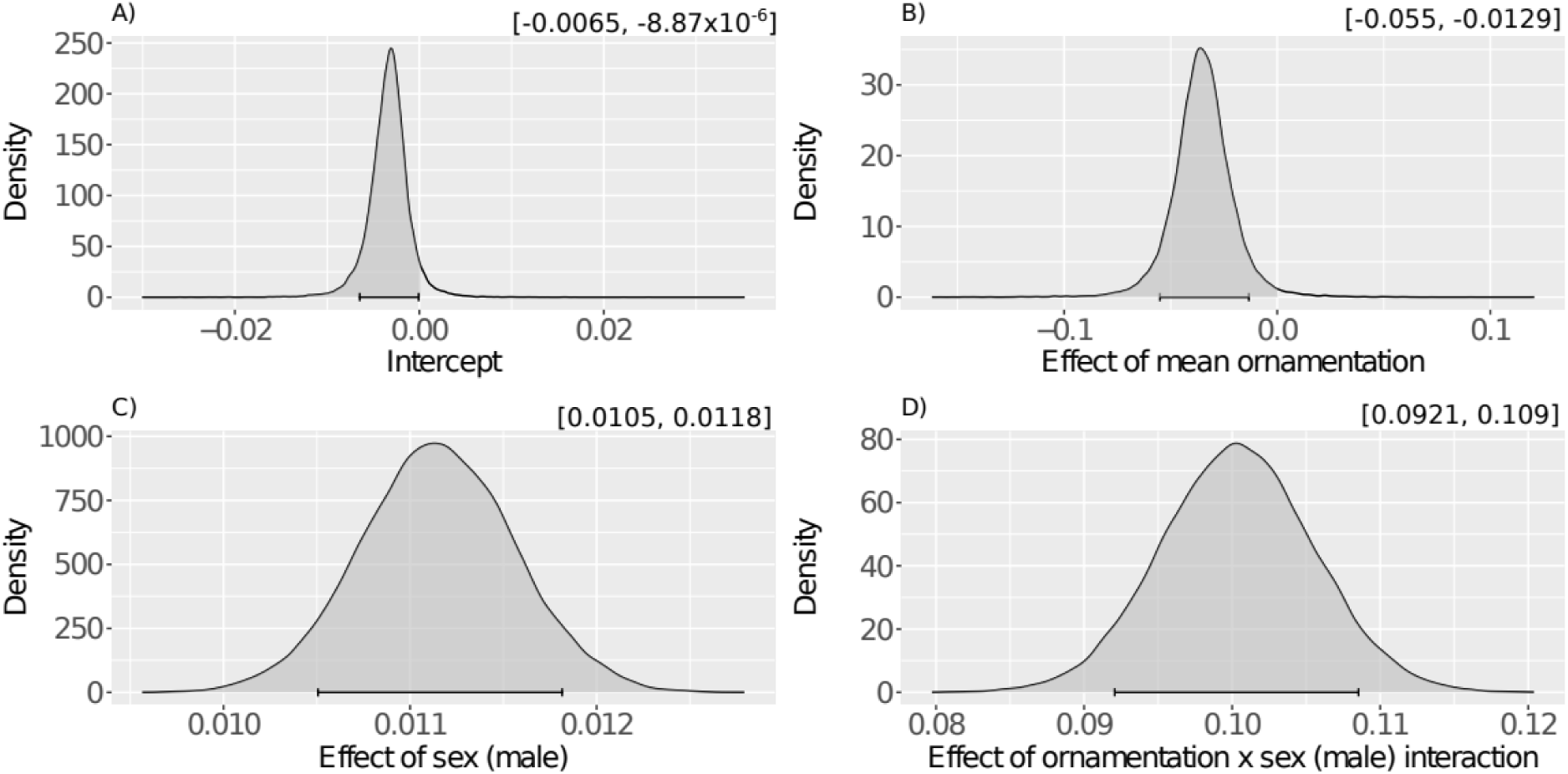
Posterior distributions of fixed effects from a phylogenetically controlled model of exaggerations of dorsal acute zones in males and females as a function of the degree of female ornamentation and sex. **A)** posterior for the intercept term, **B)** posterior for the effect of species level ornamentation, **C)** posterior for the effect of sex, and **D)** posterior for the interaction effect of species level ornamentation and sex. Shaded areas under the curve depict the proportions of the distribution < 0 (**A** and **B**) or > 0 (**C** and **D**). Horizontal error bars show the 89% HPD intervals. Inset text above each panel give the interval ranges to three significant figures. Trace plots of MCMC chains are shown in figure S7 along with traces of phylogenetic effects.

## Discussion

A number of hypotheses involving conventional sexual selection or sexual conflict have been proposed for female ornamentation and male eye exaggerations in Empidine dance flies. We show that species with more heavily ornamented females also tend to have males with more exaggerated dorsal eye facets. These findings are consistent with a process of co-evolution between these two traits.

Previous hypotheses to explain variation in male eye morphology have explicitly or implicitly invoked selection *among males* for faster mate acquisition or male mate choice (Downes 1970; Thornhill and Alcock 1983). First, eye adaptations could result from competition among males to facilitate appraisal and identification of the highest quality females. Under this scenario, exaggerated eye morphologies facilitate assessments of female quality independent of female ornamentation. However, the coincidence of both extensive female ornamentation and heightened visual ability would be surprising under this hypothesis. There would be little need for males to invest in more exaggerated eye morphologies if elaborate female ornaments were an honest signal of quality (in fact such exaggerations could be costly due to trade-offs in visual adaptations; e.g. Land [1989]). Conversely, females would have no need to invest in honest ornaments if males had heightened visual acuity to assess female reproductive value (e.g., fecundity).

Alternatively, exaggerated eye morphologies could be adaptations to “scramble-like” male competition as in other systems (e.g. Eichorn et al., 2017). However, this hypothesis predicts more exaggeration in species with strong competition among males. Because female ornaments are more prevalent in dance fly species where females are more mate-limited than males (Murray et al. 2020), this hypothesis therefore predicts a negative association between eye dimorphism and female ornament extravagance under this hypothesis. This is the opposite of our finding of positive covariance between ornament expression and male eye dimorphism (figures 3 and 4).

**Figure 4.**
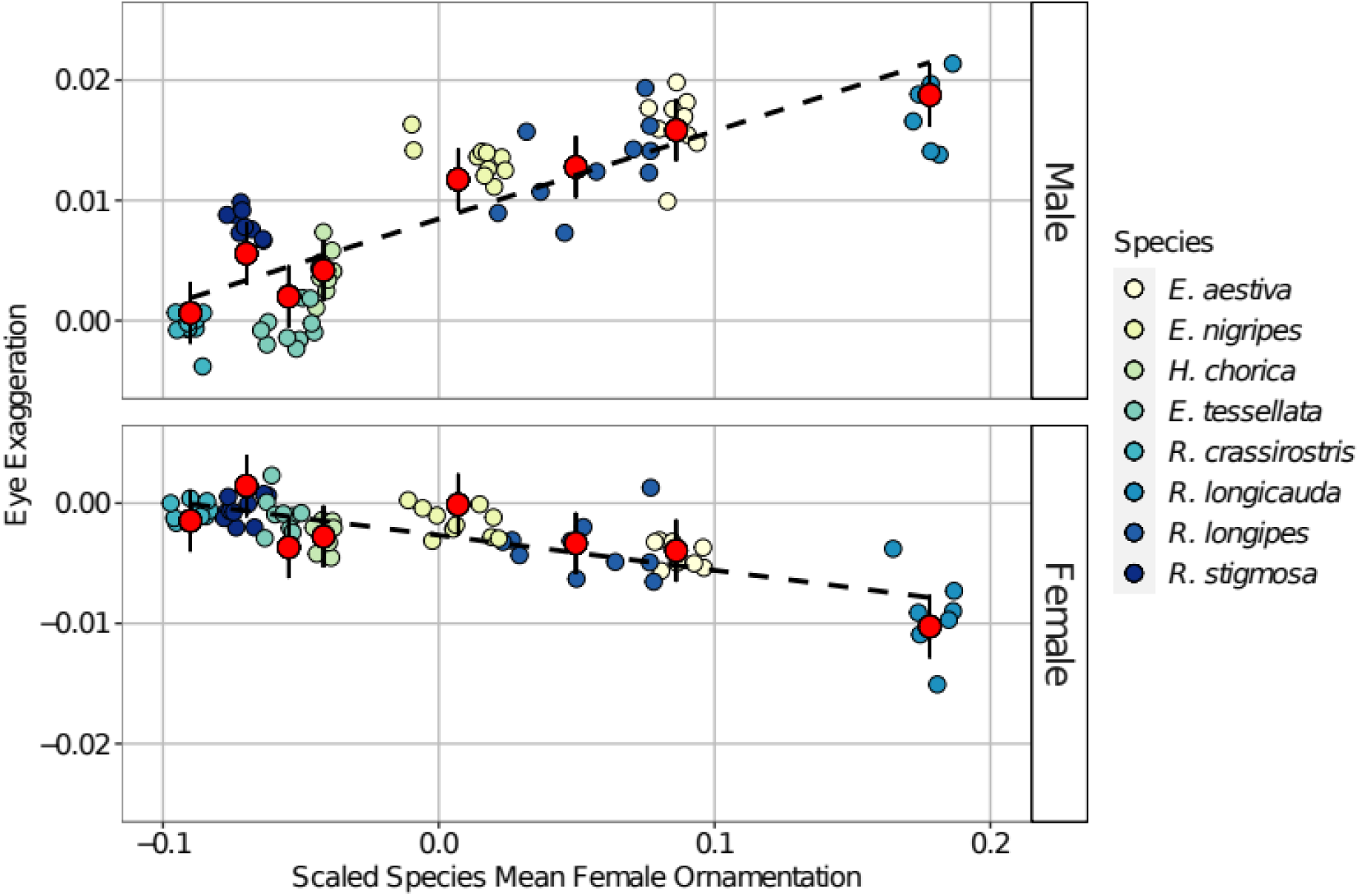
The relationship between eye exaggeration (the standardised difference between dorsal facet sizes and ventral facet sizes), for males and females, and scaled species level mean female ornamentation (as measured by standardised leg area) for 8 species of dance flies (Diptera: Empididae). Red points with error bars represent response estimates and their error from phylogenetically controlled linear models. Dashed lines represent fitted lines through the response estimates for each species and reflect the effects in figure 3. The points represent individual values for male eyes from each species. To visualise the standard error around the mean of female ornamentation for each species, we have horizontally spread points for each species in proportion to the uncertainty in species mean.

Finally, eye exaggerations might be required as part of a male preference trait for female ornaments that honestly provide signals of reproductive value (e.g. Bloch 2015). Rather than playing a role in male scramble contests, in this case the exaggerated eye morphology is the means by which males exercise preference for mates, and is expected to covary positively with the degree of female ornamentation. This pattern of positive covariation is a standard prediction of sexual selection theory where preferences and ornamental traits coevolve (Andersson 1994). However, such an association in dance flies is strange because unexaggerated eyes are undoubtedly still capable of discriminating. Eye dimorphism is not therefore a prerequisite of mate choice even if enlarged facets heighten sensitivity, and it is unclear why more conspicuous ornaments (which seem engineered to facilitate rather than hinder detection) would require the heightened sensitivity with which they cooccur. Moreover, this honest signalling hypothesis relies on female ornaments being an indicator of either direct benefits (more fecund or high condition females; LeBas et al., 2003; Browne and Gwynne 2022) or indirect benefits, such as offspring viability (good genes) or attractiveness (sexy daughters), both of which are problematic on several grounds.

Direct benefits are much more likely than indirect ones to explain male mating preferences; they can act immediately rather than awaiting expression in offspring, and do not depend on strong inter-sexual genetic correlations (Kirkpatrick and Barton 2001). Furthermore, variation in reproductive value among females should almost always exceed variation in genetic quality (Kirkpatrick and Barton 2001). In dance flies, there are two especially likely sources of variation in female quality representing direct benefits. Males might prefer females with ornaments that signal (1) greater fecundity or (2) a later stage of ovarian development (i.e., the imminence of oviposition). A female that is heavily gravid and soon to lay eggs may be especially valuable among insects with sperm storage, because such a female is less likely to mate again prior to oviposition, and remating generally decreases a focal male’s paternity by displacing his ejaculate within sperm stores (*e.g.* Boorman and Parker 1976).

However, the link between male preferences for large ornaments and ovarian status is notably inconsistent. While males sometimes prefer females with larger ornaments, the presence and intensity of sexual selection is remarkably variable across dance fly species and episodes of mating (Funk and Tallamy 2000; LeBas et al., 2003; Murray et al., 2018). In some cases there is no evidence that larger or more ornamented females acquire more mates (Hockham and Ritchie 2000; Bussière et al., 2008; Wheeler et al., 2012; Herridge 2016; Murray et al., 2019). Such inconsistency is not predicted by hypotheses of honest signalling, even if the intensity of selection can sometimes vary across populations, or through time (e.g. due to the availability of prey). Moreover, although there are usually significant and positive covariances between female ornamentation and fecundity or egg size, they can be low compared to covariances between fecundity or ovarian maturity and other indices of body size (*e.g.* thorax length; LeBas et al., 2003; Bussière et al., 2008; Murray et al., 2019; Murray et al., 2020; but see Funk and Tallamy 2000 and Wheeler 2008). In fact, regardless of the covariance between cross-sectional measures of fecundity and ornament expression, female ornaments are inherently unable to track changes in reproductive value related to ovarian development because ornaments like pinnate leg scales and wing colour are fixed at eclosion, while the number and developmental maturity of eggs changes throughout adult life (Browne and Gwynne 2022). For example, in some ornamented species, ovarian development appears to be entirely arrested prior to an initial mating and nuptial feeding event (Hunter & Bussière 2019), which means that signals of fecundity from virgin females are by necessity deceptive. Nevertheless, some recent work does suggest that female ornamentation may serve to exaggerate differences in condition, and thus provide a signal of females with greater potential fecundity, greater swarming stamina, or greater survival to oviposition (Browne and Gwynne 2022). Whether this is the case across the subfamily remains to be tested.

As an alternative to honest signalling, female ornaments in dance flies could be generally deceptive adaptations that exploit pre-existing male preferences for larger females. Deceptive ornaments would allow females to obtain nuptial gifts regardless of their current fecundity or stage of egg development (Funk and Tallamy 2000; Hockham and Ritchie 2000; Arnqvist and Rowe 2005; Arnqvist 2006), even if males prefer highly gravid females to avoid increased sperm competition risk (Herridge et al., 2016). If female ornaments are in fact deceptive signals designed to exploit male preferences for large females, we might expect resistance traits to evolve in some well-adorned species (Arnqvist and Rowe 2005). Although no resistance traits have ever been previously described among male dance flies, adaptations that augment visual perception and acuity, such as the exaggerated male eye regions observed here, are exactly the sorts of traits that are expected.

Sexual conflict and sexually antagonistic coevolution are relatively recent models proposed to explain the origin and exaggeration of sexually dimorphic traits (Parker 1979; Holland and Rice 1998; Arnqvist and Rowe 2005). Sensory exploitation of one sex by the other can play a key role in this model, with deceptive signal traits selecting for resistance in the receiver sex; the resulting arms race drives the diversification of both signal and receiver traits (Holland and Rice 1998; Arnqvist and Rowe 2005). When sensory exploitation via a seductive signal induces a suboptimal mating rate in the receiver, it increases the receiver’s exposure to costs of mating (Arnqvist 2006). To avoid costs, receivers should evolve increased discriminatory power to distinguish manipulative signals from honest ones, or to reduce susceptibility to seductive stimuli altogether. Repeated bouts of exploitation and resistance should lead to cycles of antagonistic co-evolution that can lead to a proliferation of diverse seduction and resistance adaptations (Holland and Rice 1998). Sexually antagonistic coevolution could explain seductive female traits and male resistance in rare species possessing strong male choice (Funk and Tallamy 2000; Murray et al., 2018). In such species male preference evolution should be constrained because female fecundity is often resource-limited, and males should avoid females that divert resources away from offspring towards extravagant morphologies (Fitzpatrick et al., 1995; Clutton-Brock 2009). Additionally, if females store their mates’ sperm, heavily ornamented females might acquire more mates, and consequently provide each male with a lower paternity share than less ornamented (and less polyandrous) females (Servedio and Lande 2006; Herridge et al., 2016). There is good evidence that sexually antagonistic coevolution drives diversification in traits that mediate physical mating interactions (Arnqvist and Rowe 2002; Rowe and Arnqvist 2002; Rönn et al. 2007; Bergsten and Miller 2007; Tatarnic and Cassis 2010). Sexually antagonistic sensory exploitation has also been offered as an explanation for the origins and diversification of nuptial gifts and some signal traits, and may drive the coevolution of male display traits and female behavioural responses (Sakaluk 2000; Macías Garcia and Ramirez 2005; Nakano et al. 2010; Kolm et al. 2012; Macías Garcia et al. 2012; Amcoff and Kolm 2015; Nakano et al. 2013). However, where these examples demonstrate counter adaptations in females they tend to result in dissociation between the sexual signal and the ecological signal (i.e. escape from a sexually antagonistic sensory trap), rather than co-evolving exaggerations of signal and receiver traits (e.g. Macías Garcia and Ramirez 2005; Nakano et al. 2013). To our knowledge, no previous studies have shown comparative evidence for escalating sexually antagonistic coevolution between signal and primary sensory traits, let alone in systems where females are the ornamented sex.

Sexually antagonistic coevolution, therefore, offers a compelling alternative hypothesis that overcomes several difficulties with the evolution of female ornaments, and can at once explain variation in both eye and ornamental traits across the dance flies. Female ornaments might exploit pre-existing male preferences for larger but unadorned females (because large females tend to be more gravid or more fecund; Funk and Tallamy 2000; Bonduriansky 2001; Arnqvist 2006), so that females can obtain nutritious nuptial gifts regardless of their current state of ova development. Males could then respond to female deception by developing resistance traits (increased visual perception) to avoid females with low fecundity. In the framework of sexually antagonistic coevolution, exaggerations in male eye morphology would reflect selection for increased discriminatory power to “see past” increasingly deceptive ornaments of hungry females, and identify the most gravid or fecund mates despite female disguises. This hypothesis predicts a positive association between exaggerations in eye morphology and ornamentation across species, as is seen in our data. This positive covariation would arise because exploitative adaptations that increase mating frequency for females (increased ornamentation) should be countered by increased investment in discriminatory power by males (exaggerated dorsal facet morphologies). The sexually antagonistic coevolution hypothesis also predicts a negative association between ornamentation and the proportion of males in the swarm (the swarm sex ratio) because large deceptive ornaments should be most favoured in environments where females compete more intensely for resources and experience stronger selection to deceive males (Bonduriansky 2001; Murray et al. 2020).

Moreover, our results from the literature survey suggest that exaggerations of dorsal eye morphologies in males may be specific responses to deception *via* leg pinnation. The association between ornaments and male eye exaggerations were strongest when ornamentation was defined as the presence of leg pinnation alone. However, the prevalence of pinnate scales relative to other kinds of ornaments makes strong conclusions difficult (in our sample of taxonomic descriptions, 33/93 species have females with pinnate leg scales; while only 13/93 species have other forms of female ornamentation but lack pinnation). These results hold in phylogenetically corrected analyses of a subset of these species for which we were able to collect phylogenetic information. The possibility that leg ornaments in particular select for dichoptic eyes is intriguing because the heightened photosensitivity of large eye facets may be especially effective at detecting stray photons that betray the difference between deceptively-positioned scaly legs that exaggerate abdomen size and egg-inflated large abdomens. In this context it is also noteworthy that the most extremely ornamented species, with the most exaggerated male dorsal acute zones, also have other ornamental features. *R. longicauda,* for example, has inflatable abdominal sacs that can serve to further increase a female’s apparent size to the male (Funk and Tallamy 2000; Murray *et al.,* 2018; Murray *et al.,* 2020). These inflatable sacs might also be more difficult for the male to resist by increased photosensitivity or visual acuity. In this way, additional ornamental features, such as abdominal sacs or darkened wings may represent more recent or sequential innovations in the historical arms race between seductive female characters and discriminatory male sensory structures. We also note the evidence for more exaggerated *ventral* facets among females of more ornamented species (figure 3 and figure 4). One explanation for this might be that competition among females within the swarms of species with high levels of ornamentation (and high OSR) has led to adaptations to quickly identify incoming males that typically approach swarms from below. This idea remains to be tested.

Naturally, visual perception is an important part of dance fly ecology in many other contexts apart from mate assessment. For example, hunting ability probably depends heavily on visual acuity. The sex-difference in hunting nuptial gifts could therefore conceivably lead to differences in eye dimorphism. However, there are no obvious cross-species differences in prey choice that align with variation in ornament expression or eye exaggeration (almost all empidines present nuptial gifts, including the eight species we measured quantitatively; Laurence 1952; Collin 1961; Chvala 2005). Furthermore, if hunting ability did drive the evolution of eye exaggerations, it might be strongest and most obvious in species where male competition for females is intense (i.e. high OSR, low ornamentation), to favour rapid return to mating swarms, whereas we found the converse (figures 3 and 4). We lack any evidence that hunting efficiency covaries strongly with ornament expression, and a more systematic survey of dance fly foraging ecology would help shed light on these questions.

The light environment also selects strongly on visual systems (Land 1989; 1997; Eichorn *et al.,* 2017), and some theory suggests that female ornaments are more likely in environments where male discrimination is difficult (Chenoweth *et al.,* 2006). However, no great differences in light environment (*e.g.* time of day, canopy cover) are obvious among these species from field observations (Downes 1970; Cumming 1994; Murray *et al.,* 2017; Murray *et al.,* 2018; our own observations). It is also not clear why the light environment would select for both better visual perception *and* large ornaments in females because either trait on its own would presumably improve discrimination. Nevertheless, subtle differences might play a role in adaptations toward greater visual acuity.

Our study establishes a strong signal of covariation between male eye-exaggeration and female ornamentation. We suggest that this signal is inconsistent with sexual selection in male biased swarms, and that explanations relying on honest female ornaments are problematic. We outline a plausible and intriguing alternative hypothesis involving sexual conflict and a co-evolutionary diversification of female ornaments and male visual systems. These traits provide several exciting opportunities to study the emergence of novel phenotypes, as it may be possible to use female (in the case of eye morphology) or phylogenetically “basal” morphologies as ancestral states or starting points. They also provide a novel system in which to study the inherent sexual conflicts that arise in producing sexually dimorphic phenotypes from a common genome (Pennell and Morrow 2013).

## Acknowledgements

The authors would also like to thank Adrian Plant and David Price for their invaluable assistance in species identification and natural history, NSERC, the University of Stirling, and the Scottish Centre for Ecology and the Natural Environment (SCENE) for support and accommodation during field work, and Mike Ritchie and Locke Rowe for helpful comments and discussion.

## Data and analysis code availability

All data, including phylogenetic trees, and scripts used for the statistical analyses and for producing the figures in this manuscript have been deposited in a dryad repository (https://doi.org/10.5061/dryad.rr4xgxd5z).

## Author contributions

RAWW and LFB conceived the study. RAWW, RLM, EH, VP, DTG, and LFB all contributed to the collection of specimens and data. RAWW performed the data analysis with help from RLM and LFB. RAWW led the writing of the manuscript with input from RLM, EH, DTG and LFB. All authors have approved the publication of the final manuscript.

## Ethics

Specimens of male and female Empidine dance flies were captured in the field and sacrificed by freezing until morphological measurements were made. We obtained ethical approval for all sampling and measurement procedures from the University of Stirling research ethics committee.

