## supplementary materials for "Sexually antagonistic co-evolution can explain female display signals and male sensory adaptations"

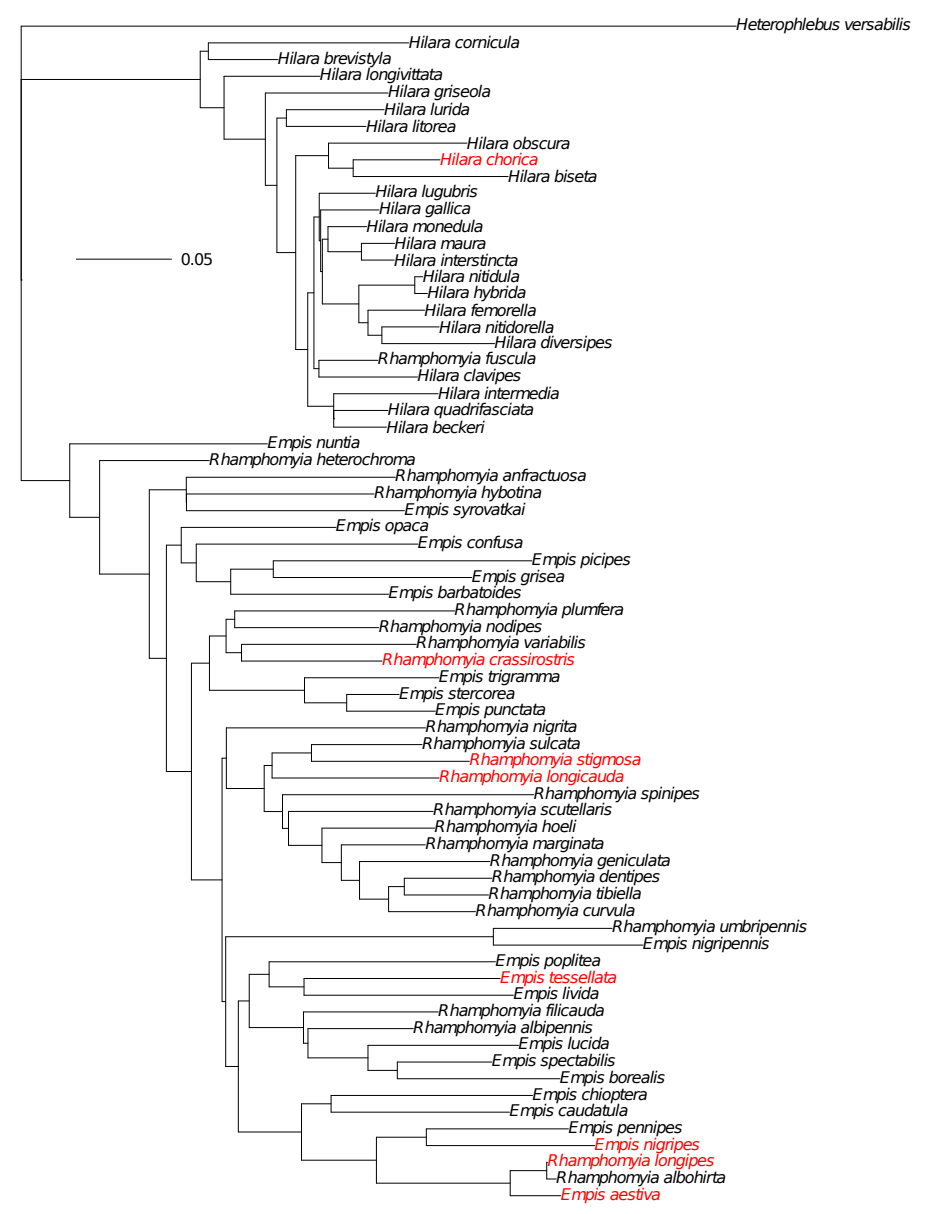
**Figure S1.** The new phylogenetic tree containing 62 species from the genera *Hilara, Empis,* and *Rhamphomyia,* as well as the outgroup *Heterophlebus versabilis.* The eight focal species for which we collected quantitative morphological data are highlighted in red. The scale bar shows the number of nucleotide substitutions per site.

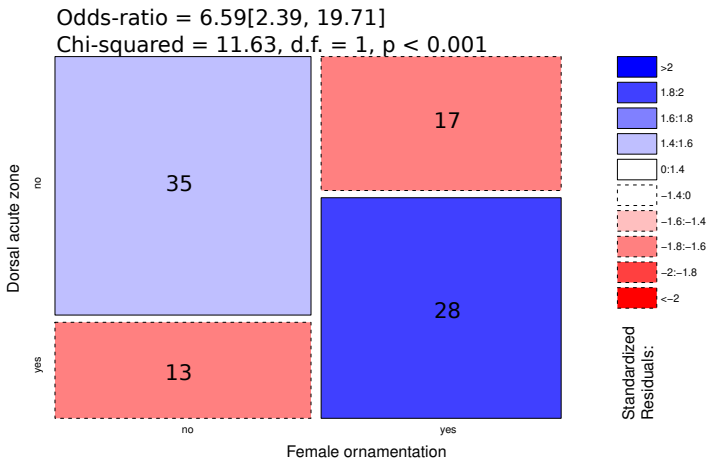
**Figure S2.** Mosaic plot of the associations between presence or absence of male dorsal acute zones and presence or absence of female ornamentation of any kind. Inset numbers give the counts of species in each area. Text at the top of the figure give relative odds of having a dorsal acute zone if the species is ornamented *versus* if the species is not ornamented, and the results of a Chi-squared test of association.

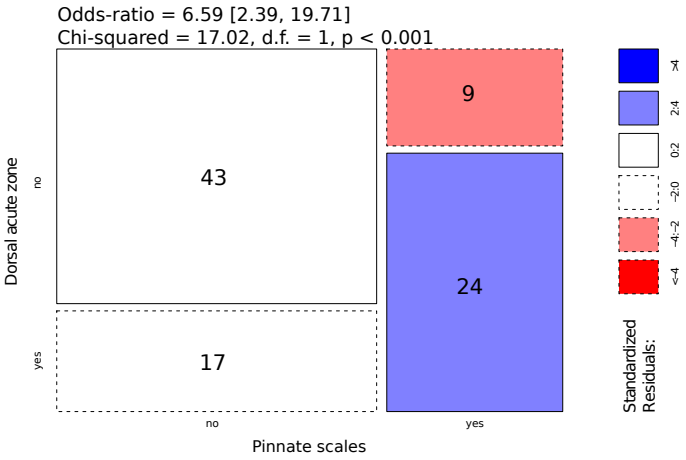
**Figure S3.** Mosaic plot of the associations between presence or absence of male dorsal acute zones and presence or absence of female pinnate scales. Inset numbers give the counts of species in each area. Text at the top of the figure give relative odds of having a dorsal acute zone if the species has pennate scales *versus* if the species does not have pennate scales, and the results of a Chi-squared test of association.

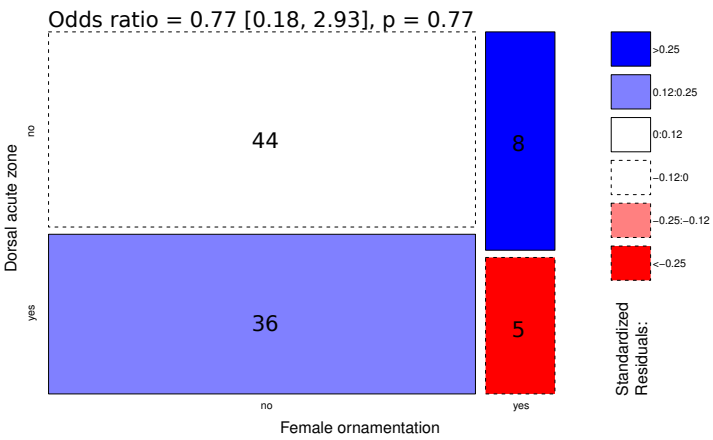
**Figure S4.** Mosaic plot of the associations between presence or absence of male dorsal acute zones and presence or absence of female ornamentation except pinnate scales. Inset numbers give the counts of species in each area. Text at the top of the figure give the results of a Chi-squared test of association.

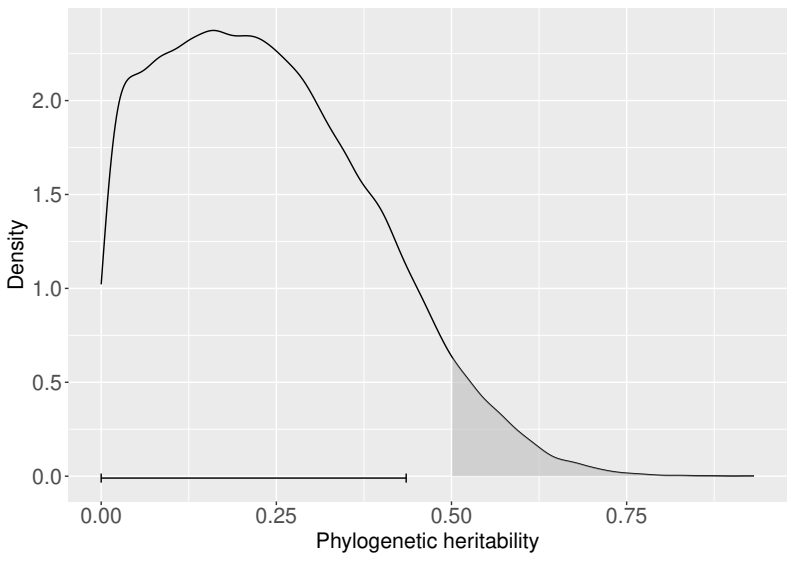
**Figure S5.** Posterior distribution of the phylogenetic heritability from a phylogenetically controlled model of exaggerations of male dorsal acute zones as a function of presence or absence of female pinnate scales. Shaded areas under the curve depict the proportions of the distribution > 0.5. Horizontal error bar shows the 89% HPD interval.

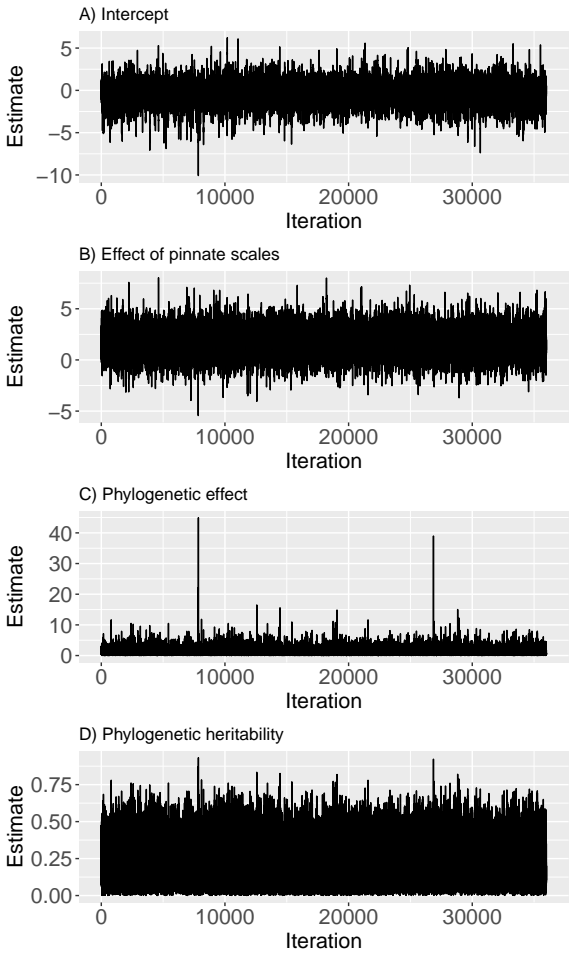
**Figure S6.** Trace plots of from a phylogenetically controlled model of exaggerations of male dorsal acute zones as a function of presence or absence of female pinnate scales. Traces are shown for **A)** the intercept, **B)** the effect of female pinnation, and **C)** the phylogenetic variance, and **D)** the phylogenetic heritability.

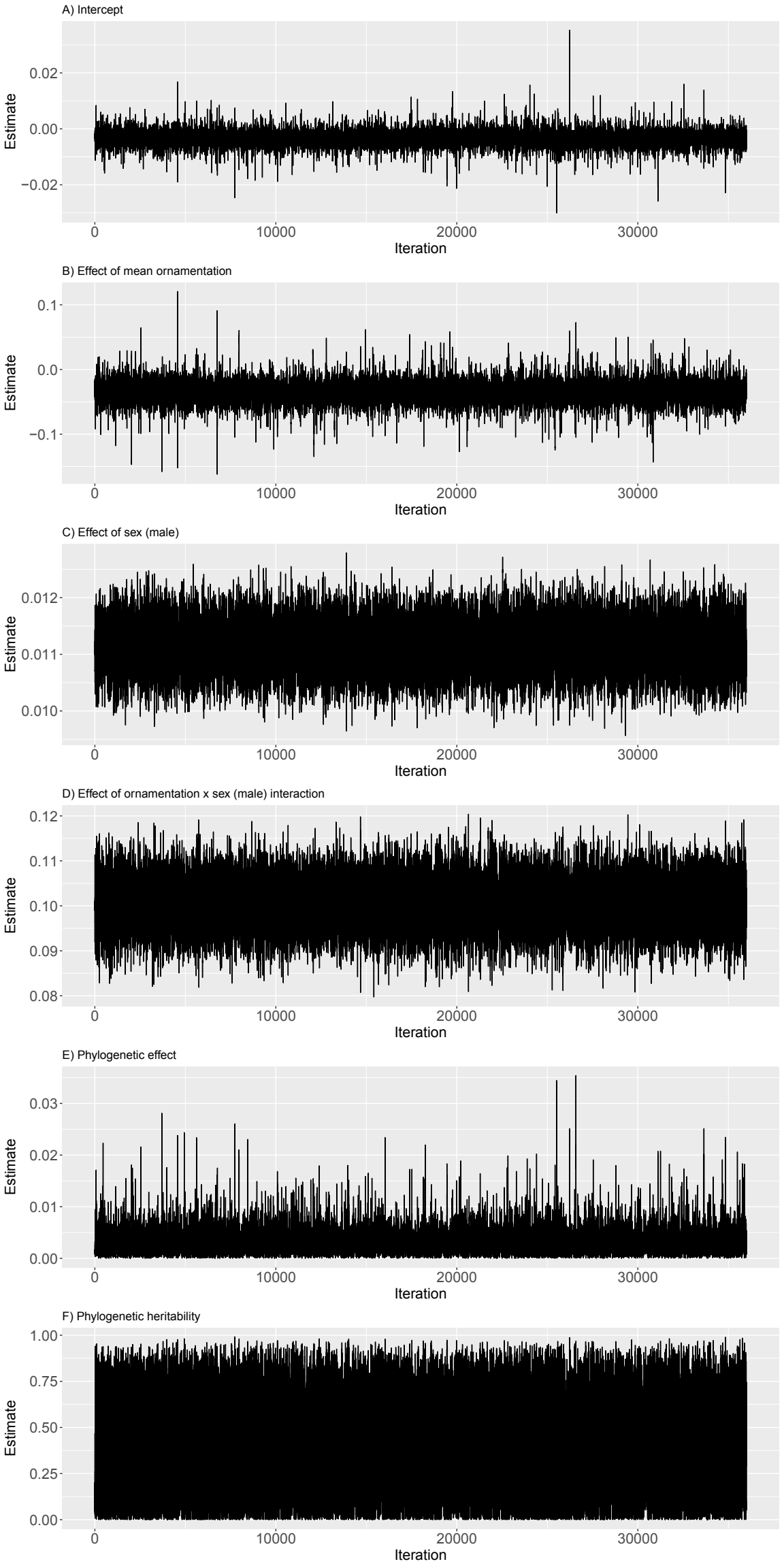
**Figure S7 (previous page).** Trace plots of MCMC chains from a phylogenetically controlled model of exaggerations of dorsal acute zones in males and females as a function of the degree of female ornamentation and sex. Traces are shown for **A)** the intercept, **B)** the partial effect of female ornamentation, **C)** the partial effect of sex (male), **D)** the effect of the sex (male) by female ornamentation interaction, and **E)** the phylogenetic variance, and **F)** the phylogenetic heritability.

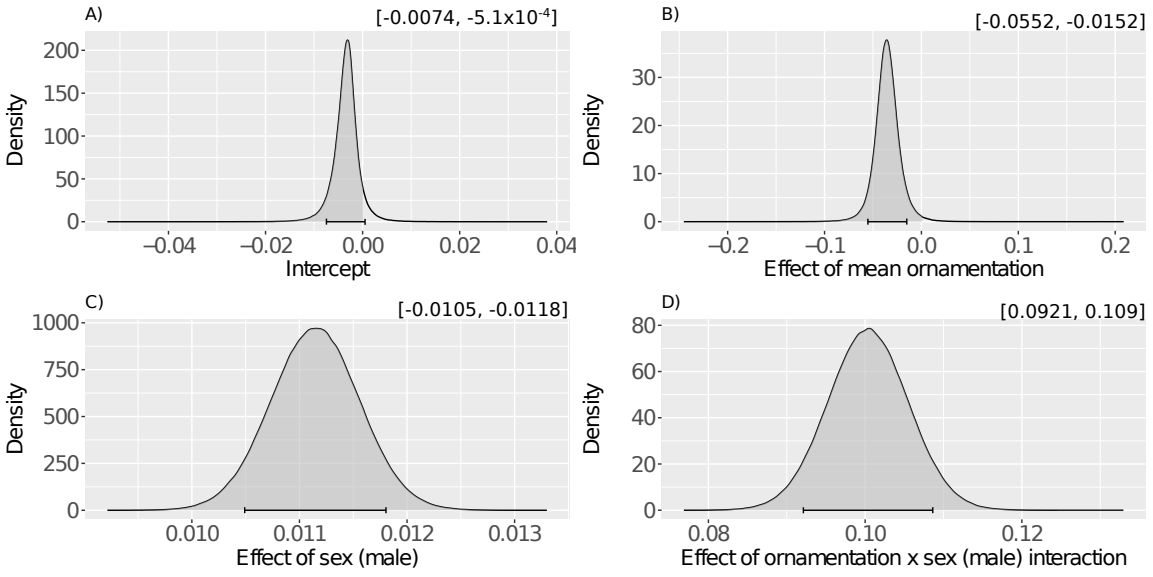
**Figure S8.** Posterior distributions of fixed effects from a phylogenetically controlled model of exaggerations of dorsal acute zones in males and females as a function of the degree of female ornamentation and sex. The model accounts for uncertainty in the tree by marginalising over a distribution of trees (see the main text for details). **A)** posterior for the intercept term, **B)** posterior for the effect of species level ornamentation, **C)** posterior for the effect of sex, and **D)** posterior for the interaction effect of species level ornamentation and sex. Shaded areas under the curve depict the proportions of the distribution < 0 (**A** and **B**) or > 0 (**C** and **D**). Horizontal error bars show the 89% HPD intervals. Inset text above each panel give the interval ranges to three significant figures. Trace plots of MCMC chains are shown in figure S8 along with traces of phylogenetic effects.

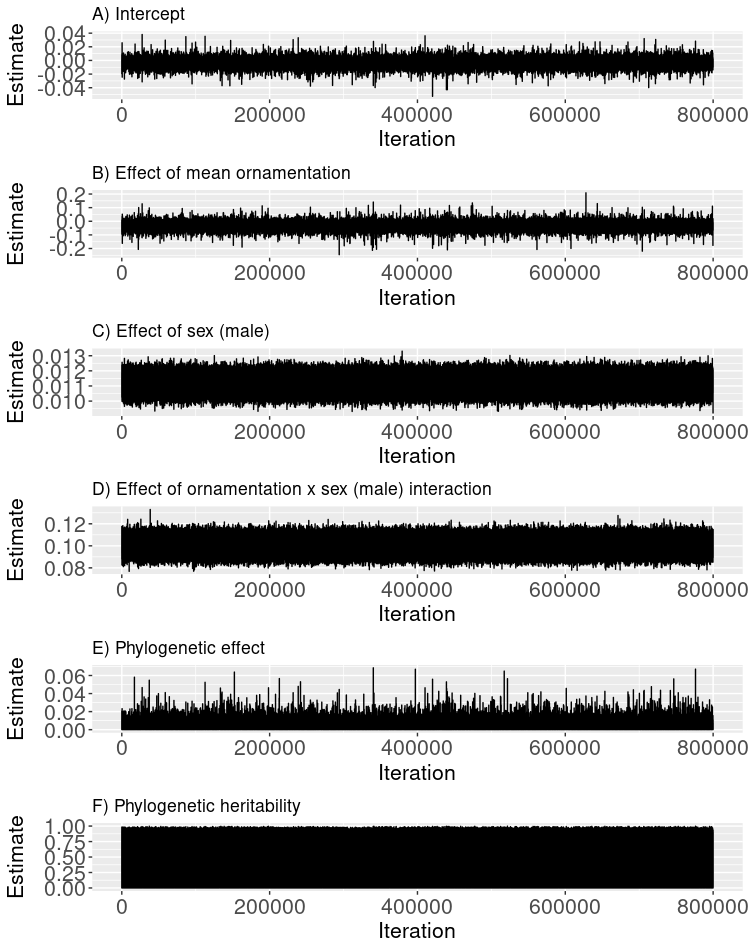
**Figure S9.** Trace plots of MCMC chains from a phylogenetically controlled model of exaggerations of dorsal acute zones in males and females as a function of the degree of female ornamentation and sex. The model accounts for uncertainty in the tree by marginalising over a distribution of trees (see the main text for details). Traces are shown for **A)** the intercept, **B)** the partial effect of female ornamentation, **C)** the partial effect of sex (male), **D)** the effect of the sex (male) by female ornamentation interaction, and **E)** the phylogenetic variance, and **F)** the phylogenetic heritability.

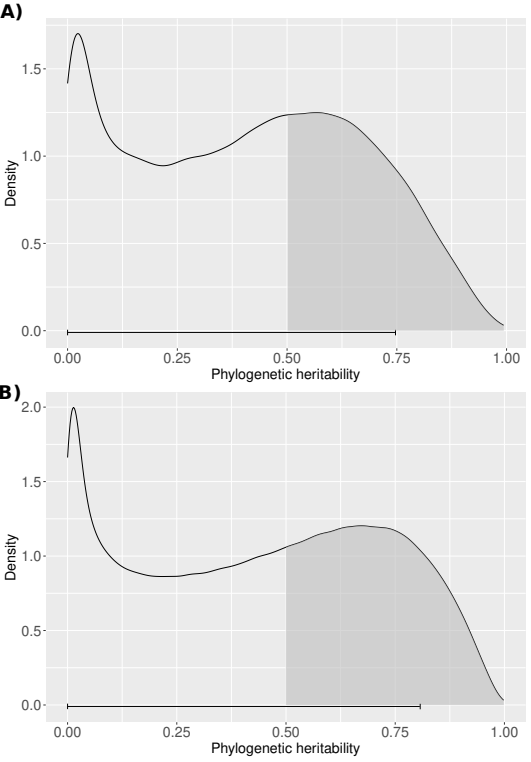
**Figure S10.** Posterior distributions of phylogenetic heritability. **A)** from a model using the consensus tree and **B)** from a model marginalising over 200 trees sampled from the posterior distribution of trees. Horizontal error bars show the 89% HPD intervals.

**Table S1.** Summary statistics for each species, including the mean and standard error (SE) for several morphological traits measured in males (M) and females (F). Thorax length is given in original units. Male and female eye exaggeration are standardised by thorax length. Ornamentation values are provided for all females sampled for each species, and as such are values prior to scaling (see Methods in the main text).

|  | ***N*** | | ***Ornamentation [SE]*** | ***Eye exaggeration***  ***[SE]*** | | ***Thorax length (mm)***  ***[SE]*** | |
| --- | --- | --- | --- | --- | --- | --- | --- |
| ***Species*** | ***M*** | ***F*** | ***F*** | ***M*** | ***F*** | ***M*** | ***F*** |
| *R. longicauda* | 6 | 7 | 0.26  [0.014] | 0.017  [0.0013] | -0.0093 [0.0013] | 1.35  [0.061] | 1.38  [0.083] |
| *R. longipes* | 10 | 10 | 0.14  [0.029] | 0.013  [0.0011] | -0.0037  [0.00072] | 0.85  [0.02] | 0.86  [0.018] |
| *R. stigmosa* | 9 | 10 | 0.017  [0.0082] | 0.008  [0.00037] | -0.00056  [0.00032] | 1.71  [0.031] | 1.67  [0.034] |
| *R. crassirostris* | 10 | 10 | -0.0037  [0.0073] | -0.0005  [0.00041] | -0.00056  [0.00021] | 1.8  [0.046] | 1.79  [0.052] |
| *E. aestiva* | 8 | 9 | 0.172  [0.011] | 0.016  [0.00094] | -0.0043  [0.00038] | 0.92  [0.023] | 0.99  [0.044] |
| *E. tessellata* | 9 | 9 | 0.045  [0.0056] | 0.004  [0.00061] | -0.0027  [0.00038] | 3.55  [0.078] | 3.33  [0.086] |
| *E. nigripes* | 10 | 10 | 0.093  [0.021] | 0.013  [0.00045] | -0.0015  [0.00039] | 1.13  [0.025] | 1.11  [0.018] |
| *H. chorica* | 10 | 9 | 0.032  [0.011] | -0.0006  [0.00046] | -0.00097  [0.00051] | 0.85  [0.041] | 0.86  [0.014] |
